# T2T genomes of *Caenorhabditis nigoni* and *Caenorhabditis briggsae* reveals extensive loss of satellite DNA associated with self-fertilization

**DOI:** 10.64898/2026.02.11.705409

**Authors:** Ryan Pellow, Manitejus Kotikalapudi, Ofer Rog

## Abstract

The two closely related *Caenorhabditis* nematode species, *C. nigoni* and *C. briggsae*, are commonly used to study the evolution of reproductive modes in animals, with the self-fertile *C. briggsae* and outcrossing *C. nigoni* sharing a common ancestor ∼3.5 million years ago. Earlier genomic analyses of these species revealed genome shrinkage associated with selfing and proposed that at least some gene loss can be adaptive. However, the incomplete *C. nigoni* reference genome limited most comparative analyses to genic regions. Here, we leveraged long-read sequencing to generate a telomere-to-telomere (T2T) assembly for the *C. nigoni* strain JU1422 and the *C. briggsae* strain AF16. This new 139Mb *C. nigoni* genome resolved 57 gaps and 149 unassigned scaffolds from the previous genome assembly. Comparison with the 107Mb T2T *C. briggsae* genome reveals that the major driver of genome content differences are deletions to satellite DNA arrays, reflecting a loss of 9.6Mb. Interestingly, many of the differences are on the *C. nigoni* X chromosome, which is >13Mb larger than in the previous assembly. The transition to selfing was thus accompanied by a 37% reduction in the size of the sex chromosome compared to 16-21% shrinkage of the autosomes. We also document a surprising degree of plasticity in the ribosomal DNA, with the X chromosome harboring a second 45S rDNA array that is absent in *C. briggsae*. Our analysis reveals that obligatory outcrossing may play a major role in the maintenance of satellite DNA arrays.

## Introduction

High-quality reference genomes facilitate downstream genomic analyses and provide the means to study the relationships between genotypes and phenotypes. The nematode *Caenorhabditis elegans* is a key model organism in biology and in 1998, was the first animal to have its genome sequenced (Consortium* 1998). By 2005, its genome was considered complete and gapless (Hillier et al. 2005), but only recently, by leveraging advances in long-read technologies, has a gap-less, telomere-to-telomere (T2T) genome been assembled (Tyson et al. 2018; Yoshimura et al. 2019; Ichikawa et al. 2025). Among many findings, the new assembly revealed many hitherto unknown repetitive regions (satellite DNA; satDNA) that comprise 6.6% of the genome.

*Caenorhabditis* has served as great model to study genome evolution, with analysis within the Elegans supergroup proving particularly fruitful. *C. briggsae* and *C. nigoni* diverged from each other ∼3.5 million years ago (Thomas et al. 2015) and diverged from *C. elegans* ∼100 million years ago (Coghlan and Wolfe 2002). Interestingly, the higher-order structure of the genomes in *Caenorhabditis* is generally conserved, with each species harboring six holocentric chromosomes and relatively few genes translocated between chromosomes (Albertson and Thomson 1982; Hillier et al. 2007; Zedek and Bures 2012; Carlton et al. 2022; Caro et al. 2022). This conservation in chromosome organization contrasts with high levels of divergence at the nucleotide level and some of the highest rates of genomic rearrangements in the animal kingdom (4.1-9.7 genomic rearrangements per million years (Coghlan and Wolfe 2002; Hillier et al. 2007; Wang et al. 2025)).

Within *Caenorhabditis, C. briggsae* and *C. nigoni* are the most closely related and exhibit interfertility (Woodruff et al. 2010; Kozlowska et al. 2012; Felix et al. 2014; Ryan and Haag 2017; Devi et al. 2025). Due to this, they have been used to study hybrid incompatibility and the evolution of reproductive modes in animals (reviewed in (Thomas et al. 2012b; Cutter 2018; Cutter et al. 2019)). More specifically, *C. briggsae* is an androdiecious species, meaning its primary mode of reproduction is self-fertilization via hermaphrodites, with rare males allowing for gene flow between populations. *C. nigoni* is gonochoristic and reproduces exclusively through outcrossing. Self-fertilization benefits from reproductive assurance and is thought to allow efficient dispersion and rapid colonization of new niches, which compensates for the loss of genetic diversity (Thomas et al. 2012b; Theologidis et al. 2014). This point is highlighted by *C. briggsae* having an effective population size that is orders of magnitude lower than its outcrossing counterparts (Teterina et al. 2025). This is despite *C. briggsae* having a wide, cosmopolitan global distribution, whereas *C. nigoni* is restricted to tropical regions (Moya et al. 2025). Both species, as all other studied *Caenorhabditis* species, possesses an *XO* sex determination system where *XO* individuals are male, and *XX* individuals are either female or self-fertilizing hermaphrodites, depending on the species.

Self-fertile hermaphrodites have evolved independently three times in *Caenorhabditis* (*C. briggsae*, *C. elegans*, and *C. tropicalis*; reviewed in (Ellis 2017)) (Hill et al. 2006; Guo et al. 2009; Hill and Haag 2009; Kiontke et al. 2011; Chen et al. 2014). These transitions are associated with a 10-30% reduction in genome size relative to their most closely related obligatory outcrossing species (Thomas et al. 2012a; Fierst et al. 2015; Yin et al. 2018; Adams et al. 2023). However, a mechanism to explain this observation remains unclear as current hypotheses include both genome shrinkage (Thomas et al. 2012a; Fierst et al. 2015; Yin et al. 2018) and genome growth models (Kanzaki et al. 2018; Adams et al. 2023). Comparative genomics studies revealed that *C. nigoni* contains more protein-coding genes than *C. briggsae,* and that the genes absent from the *C. briggsae* genome are disproportionately male-biased in expression, linking at least some of this loss to altered evolutionary pressures brought about by self-fertilization (Yin et al. 2018; Adams et al. 2023). More broadly, while genome shrinkage has been associated with transition to self-fertilization in plants, the universal effects of self-fertilization, if any, and the mechanisms underlying them remain a matter of debate. A major caveat to these previous analyses is the lack of high-quality, complete reference genomes, limiting most analyses to genic regions.

Here, we generated T2T reference genomes for the *C. nigoni* strain JU1422 and the *C. briggsae* strain AF16 using Nanopore sequencing. The substantially improved *C. nigoni* reference genome allows us to attribute previously unassigned and unaccounted for sequences, including a second 45S rDNA array located on the X chromosome. We revisit comparative genomics analyses with regards to genome content, genome size, and genome rearrangements to reveal that the genome of *C. briggsae* experienced large-scale deletions of satDNA, especially on the X chromosome, and high rates of large inversions.

## Results

### Genome sequencing and T2T assemblies

Of the two species, *C. briggsae* has a substantially more complete reference genome. This is due to *C. briggsae* being widely used as a satellite model organism to *C. elegans* (Baird and Chamberlin 2006; Cutter 2015; Stevens et al. 2022), and the ease of obtaining inbred strains via self-fertilization. The first reference genome of *C. briggsae* used the strain AF16 and was generated in 2003, just five years after the publishing of the *C. elegans* reference genome (Consortium* 1998). This reference genome was constructed using Sanger-based shotgun sequencing and a physical map (Stein et al. 2003). It was further refined following the construction of genetic maps (Hillier et al. 2007; Ross et al. 2011) and later by combining multiple sequencing methods (Nanopore long-read, Illumina short-read, and Hi-C) to generate a reference genome containing less than 10 gaps (Stevens et al. 2022). In contrast, much lower quality reference genomes exist for the *C. nigoni* strains JU1421 (Ren et al. 2018) and JU1422 (Yin et al. 2018). Both strains are the result of at least 25 generations of inbreeding and derived from the “population strain” JU1325, which represents a collection of individuals obtained from the wild (Kozlowska et al. 2012). For this study, we chose to generate T2T reference genomes for the *C. briggsae* strain AF16 and the *C. nigoni* strain JU1422 because of their previous use as reference strains and their ubiquitous use in comparative genomics studies. We confirmed some of our findings by analyzing sequencing reads from a second *C. nigoni* strain (EG5268), which is highly diverged from JU1422 (Kiontke et al. 2011).

To generate T2T reference genomes for *C. briggsae* AF16 and the *C. nigoni* JU1422, we sequenced each strain to over 40x coverage using the Oxford Nanopore MinION platform (**Supplemental Table 1**). Although the read coverage is moderate compared to previous Nanopore-based reference assemblies (219x for *C. briggsae* (Stevens et al. 2022) and 96x for *C. nigoni* (Yin et al. 2018)), our mean read lengths were substantially longer (28,014bp and 28,535bp compared to 15,705 and 8,469bp for *C. briggsae* and *C. nigoni*, respectively). Particularly important metrics for the assembly were N50 read lengths close to 50kb (48,242bp (*C. briggsae*) and 47,593bp (*C. nigoni*)) and over 5x coverage with reads >100kb (7.3x and 5.8x for *C. briggsae* and *C. nigoni*, respectively)

To convert these high-quality reads to T2T reference genomes, initial assemblies were generated using Flye (Kolmogorov et al. 2019). However, these initial Flye assemblies still contained incorrectly assembled regions and gaps, as well as incorrect bases stemming from non-random, strand-specific basecalling errors (total of 82 and 66 for *C. briggsae* and *C. nigoni*, respectively; **Supplemental Figures 1-3**). These gaps and issues were manually resolved (similar to (Ichikawa et al. 2025)) by using a combination of Winnowmap2 (Jain et al. 2022) and IGV (Thorvaldsdottir et al. 2013), while the basecalling issues were remedied by polishing the affected regions with publicly available PacBio or Illumina reads (Albritton et al. 2014; Yin et al. 2018). In all cases, correct assembly was confirmed by remapping of sequencing reads and manual inspection of mismatches and genome coverage (see Methods).

The resulting T2T genomes for *C. briggsae* and *C. nigoni* were 107.1Mb and 139.0Mb, respectively, and represent a 1.9Mb and 21.2Mb increase over previous assemblies (**Table 1**; the increase over the total size, including previously unassigned scaffolds is 1.6Mb and 9.6Mb, respectively). Importantly, the BUSCO completeness scores (capturing the presence and number of common essential single-copy genes; (Simao et al. 2015)) and the total number genes predicted by the gene prediction tool Augustus (Stanke et al. 2006) remain similar in both the old and new references, consistent with the idea that non-genic regions are responsible for most of the gaps in previous reference genomes. While the genome size of this new *C. briggsae* AF16 reference genome is in line with recent, mostly complete genome assemblies, the genome size of this new *C. nigoni* JU1422 reference genome is substantially more complete than the previous reference genome (Yin et al. 2018), resolving 58 gaps and 149 unassigned scaffolds (**Figure 1**). Below, we analyze the salient features revealed by these complete, gapless T2T reference genomes of *C. briggsae* and *C. nigoni*.

**Figure 1:**
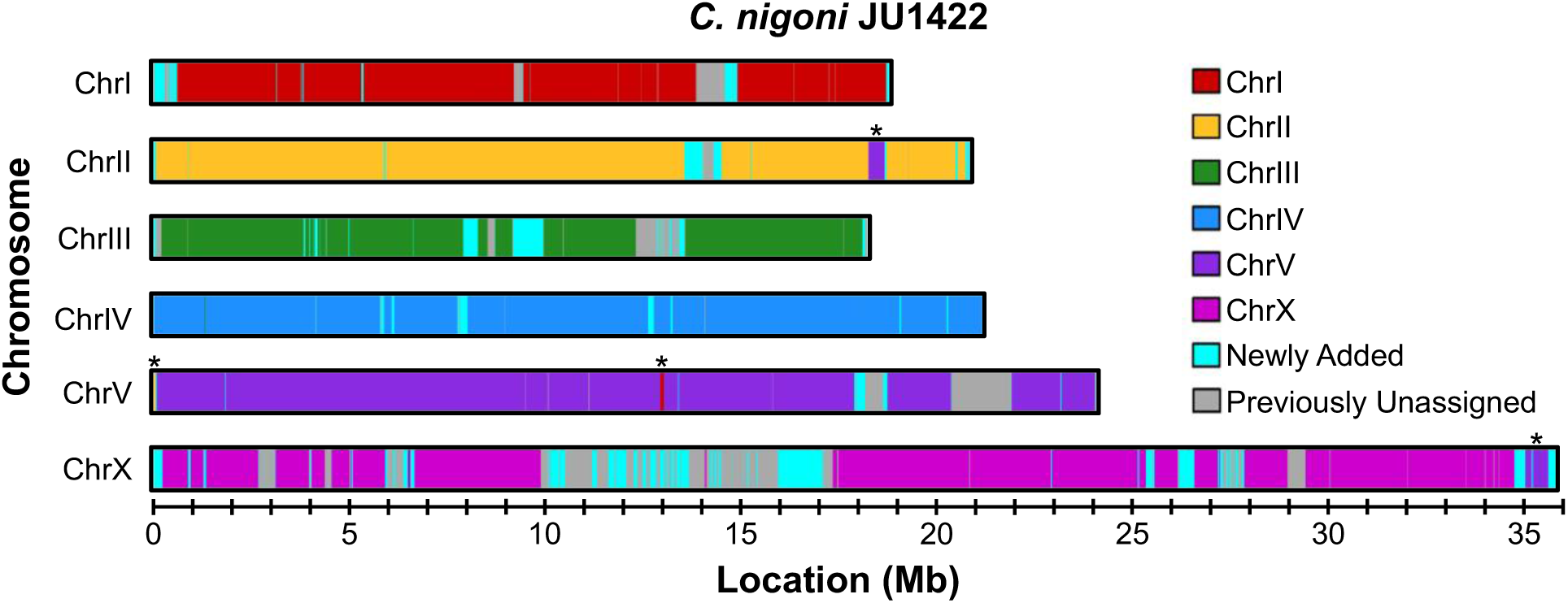
Chromatograms depicting the T2T assembly of the *C. nigoni* JU1422 genome. The colors red, yellow, green, blue, purple, and pink represent the sequences and chromosome assignments from the previous *C. nigoni* JU1422 reference genome. Cyan indicates sequences that were not present in the scaffolds of the previous assembly, while grey denotes sequences that were present as scaffolds in the previous assembly but not assigned to a chromosome. Asterisks denote previously incorrectly assigned segments on chromosomes II, V and X.

**Table 1.** **Assembly statistics for *C.briggsae* and *C. nigoni***

|  | *C. briggsae* AF16 | *C. briggsae* AF16 | *C. briggsae* QX1410 | *C. briggsae* VX34 | *C. nigoni* JU1422 | *C. nigoni* JU1422 |
| --- | --- | --- | --- | --- | --- | --- |
| Reference | This Study | Ross et al. 2011 | Stevens et al. 2022 | Stevens et al. 2022 | This Study | Yin et al. 2018 |
| Total Size | 107,058,359 | 108,384,165 | 106,196,058 | 106,953,292 | 139,019,943 | 129,488,540 |
| Total Size w/o N's | 107,058,359 | 105,418,665 | 106,192,558 | 106,951,792 | 139,019,943 | 129,435,387 |
| Span of N's | 0 | 2,965,500 | 3,500 | 1,500 | 0 | 53,153 |
| Scaffolds | 6(+MT) | 367 | 6(+MT) | 6(+MT) | 6(+MT) | 155 |
| Unassigned Scaffolds | 0 | 361 | 0 | 0 | 0 | 149 |
| Contigs | 6(+MT) | 5,074 | 14 | 10 | 6(+MT) | 213 |
| Size of Six Chromosome Scaffolds | 107,043,955 | 105,183,150 | 106,181,464 | 106,938,796 | 139,005,504 | 117,831,421 |
| Gaps in Six Chromosome Scaffolds | 0 | 4,707 | 7 | 3 | 0 | 58 |
| BUSCO | 99.33 | 99.33 | 99.33 | 99.33 | 99.16 | 99.16 |
| Protein-Coding Genes on Six Scaffolds | 25,171 | 24,635 | 24,983 | 25,119 | 33,148 | 29,728* |
| * Protein coding genes on all scaffolds = 32,575 |  |  |  |  |  |  |

### Rapid shrinkage of the *C. briggsae* genome is associated with the loss of satDNA

Across the Elegans supergroup within the *Caenorhabditis* genus, self-fertile hermaphrodite species have arisen at least three times, with each transition associated with a 10-30% reduction in genome size (Thomas et al. 2012a; Fierst et al. 2015; Yin et al. 2018; Adams et al. 2023). Due to each transition occurring independently, the genome size of the outcrossing species (in our case, *C. nigoni*) is inferred to be closer to the ancestral genome size.

We sought to revisit the differences in genome size between *C. briggsae* and *C. nigoni* using our significantly improved genomes. The *C. nigoni* reference genome is now 21.2Mb larger than previous genome size estimates and 29.9% larger than *C. briggsae* (**Supplemental Table S2**). This updated genome size aligns with the genome sizes of the more distantly related outcrossing species (Fierst et al. 2015; Yin et al. 2018; Stevens et al. 2019). Interestingly, we find that majority (57.4%) of the newly assembled sequences belong to the X chromosome. This results in the size of the X chromosome increasing by 51.4% relative to previous assemblies.

This X-specific effect was also observed when comparing the new assemblies on the chromosomal level. We find that the *C. briggsae* autosomes have experienced 15.2-20.9% reductions in size relative to *C. nigoni*, while the X chromosome has shrunk by 37.4% (**Table 2**).

**Tabel 2.** Chromosome length comparison between *C. nigoni* JU1422 and *C. briggsae* AF16.

| Table 2 Chromosome length comparison between *C. nigoni* JU1422 and *C. briggsae* AF16 |  |  |  |  |
| --- | --- | --- | --- | --- |
| Chromosome | JU1422 Size (bp) | AF16 Size (bp) | Difference | Percent |
| ChrI | 18,859,249 | 15,453,935 | -3,405,314 | -18.1% |
| ChrII | 20,855,734 | 16,504,556 | -4,351,178 | -20.9% |
| ChrIII | 18,237,684 | 14,859,658 | -3,378,026 | -18.5% |
| ChrIV | 21,159,230 | 17,358,786 | -3,800,444 | -18.0% |
| ChrV | 24,086,101 | 20,434,183 | -3,651,918 | -15.2% |
| ChrX | 35,807,506 | 22,432,837 | -13,374,669 | -37.4% |
| Mitochondria | 14,439 | 14,404 | -35 | -0.2% |
| **Total** | **139,019,943** | **107,058,359** | **-31,961,584** | **-23.0%** |

To validate this drastic and previously unappreciated reduction in the size of the X chromosome, we imaged the germline of the two nematodes. We identified the X chromosome based on the absence of histone H3 lysine 36 trimethylation (H3K36me3), which is enriched on autosomes (Kelly et al. 2002; Bender et al. 2006; Rechtsteiner et al. 2010; Gao et al. 2015) (**Figure 2A-B**). By measuring the DAPI intensity of the X chromosome and comparing it to the total DAPI intensity, we find that the median DAPI intensity of the X chromosome accounts for 19.7% of the *C. briggsae* genome and 26.0% for the *C. nigoni* genome, in agreement with the fraction of the X chromosomes in our genome assemblies: 21.0% (*C. briggsae*) and 25.8% (*C. nigoni*) (**Figure 2C**) (**Supplemental Table S3**).

**Figure 2:**
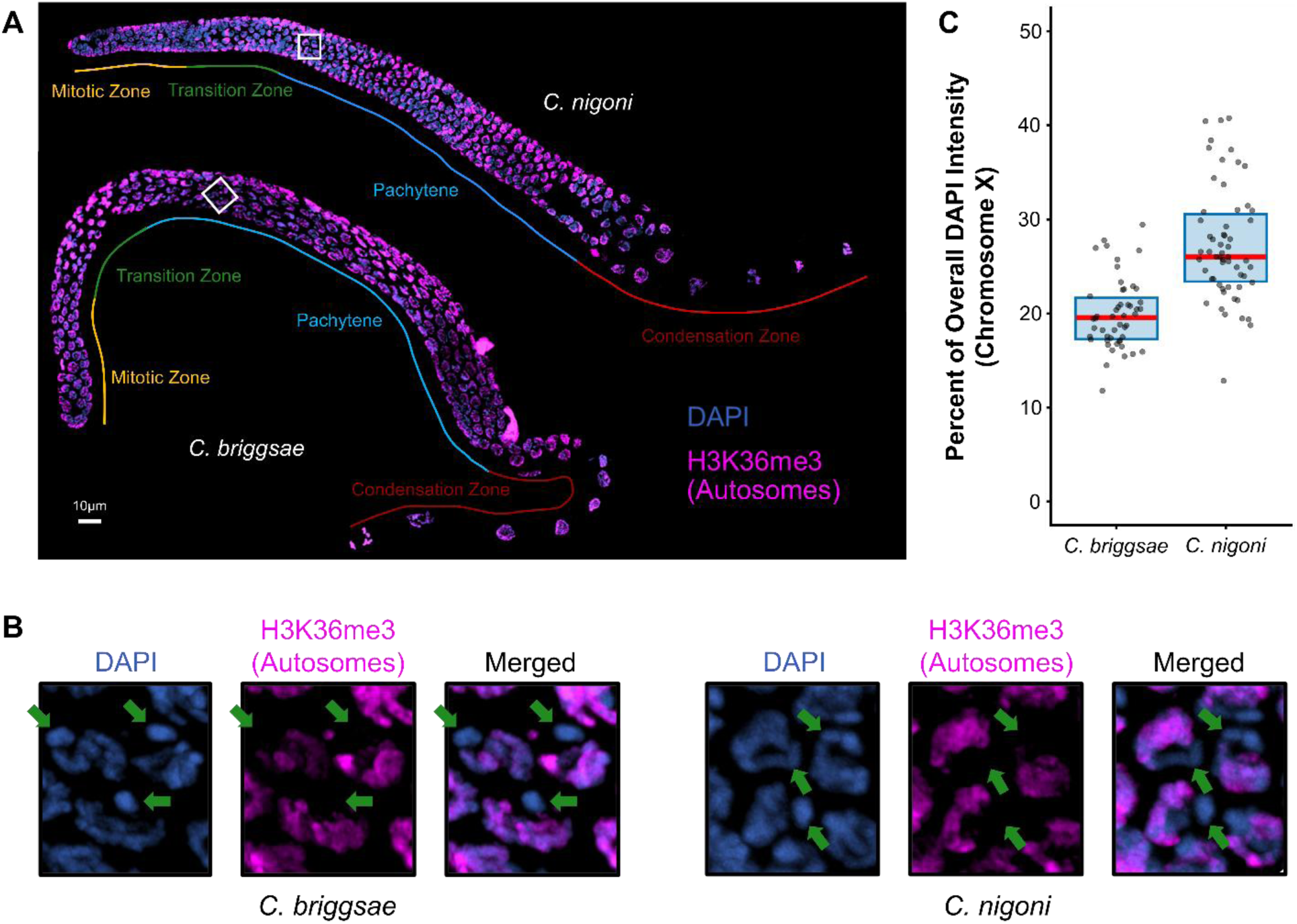
Cytological comparison between *C. briggsae* and *C. nigoni* X chromosomes. (**A**) Maximum intensity projection of immunofluorescence images of *C. briggsae* and *C. nigoni* oogenic germlines. H3K36me3 is shown in pink. DAPI is shown in blue. Meiotic progression is labelled as in (Shakes et al. 2009). (**B**) Zoomed-in views of early pachytene nuclei of *C. briggsae* (left) and *C. nigoni* (right). Green arrows highlight the X chromosome which, unlike the autosomes, lacks the methylation mark H3K36me3. (**C**) Boxplots depicting the percentage of overall DAPI intensity that is accounted for by the X chromosome in *C. briggsae* and *C. nigoni*. Red line indicates the median, and the box indicates represents the 25^th^ and 75^th^ quartiles. Each datapoint represents a single nucleus. N=50 and N=58 nuclei for *C. briggsae* and *C. nigoni*, respectively. Three gonads from each species were analyzed.

To investigate genetic patterns accounting for the reduced *C. briggsae* genome size, we compared the presence of local homology as well as large-scale synteny between *C. briggsae* and *C. nigoni*. Homology was determined by probing local alignment, using BLAST, of consecutive 10kb windows from the *C. nigoni* genome against the homologous *C. briggsae* chromosome and vice versa (**Supplemental Figure S4**). By treating each 10kb window independently, this method allows to distinguish large duplications in the *C. nigoni* lineage from large deletion in the *C. briggsae* lineage. We observed the presence of large regions in the *C. nigoni* genome that lack a homologous sequence in *C. briggsae*, as well as a relative paucity of *C. briggsae* regions that lack a homologous sequence in *C. nigoni*. This pattern is consistent with contraction of the *C. briggsae* genome rather than expansion of the *C. nigoni* genome (see **Methods** and **Discussion**).

In the autosomes, the chromosome centers exhibit more homology than the chromosome arms, which is consistent with lower intraspecies single-nucleotide polymorphism (SNP) distribution, infrequent meiotic recombination, and fewer hyper-divergent regions (Rockman and Kruglyak 2009; Ross et al. 2011; Lee et al. 2021; Noble et al. 2021; Moya et al. 2025). On the X chromosome, non-homologous regions were present throughout the chromosome, including large regions in the chromosome center where an almost contiguous block >8Mb share little to no identity with the *C. briggsae* counterpart. Synteny analysis using global alignment confirmed these patterns. We tested the ability to align the *C. briggsae* genome to the *C. nigoni* genome, and defined regions where alignment was identified as ’conserved’, while gaps in alignment were defined as ’diverged’. As expected, the two approaches were significantly correlated, with most diverged regions appearing on the arms of autosomes and throughout the X chromosomes (Pearson Correlation Coefficient (*r*) = 38.6, *p*-value < 1 x 10^-308^) (**Figure 3A**). As above, the inverse global alignment - of the *C. nigoni* genome to *C. briggsae* - revealed fewer and smaller diverged regions. Importantly, 82.2% of the base pairs lost in *C. briggsae* are in regions that are diverged between the two species, with the diverged regions on the X chromosome responsible for the largest proportion of lost sequence (**Figure 3B and 3C**) (**Supplemental Table S4**). This is despite the diverged regions only accounting for 25.7% of the *C. nigoni* genome. Overall, the patterning of conserved and diverged regions implies that the genome shrinkage that happened as *C. briggsae* transitioned to androdioecy, occurred as large deletions rather than large duplications.

**Figure 3:**
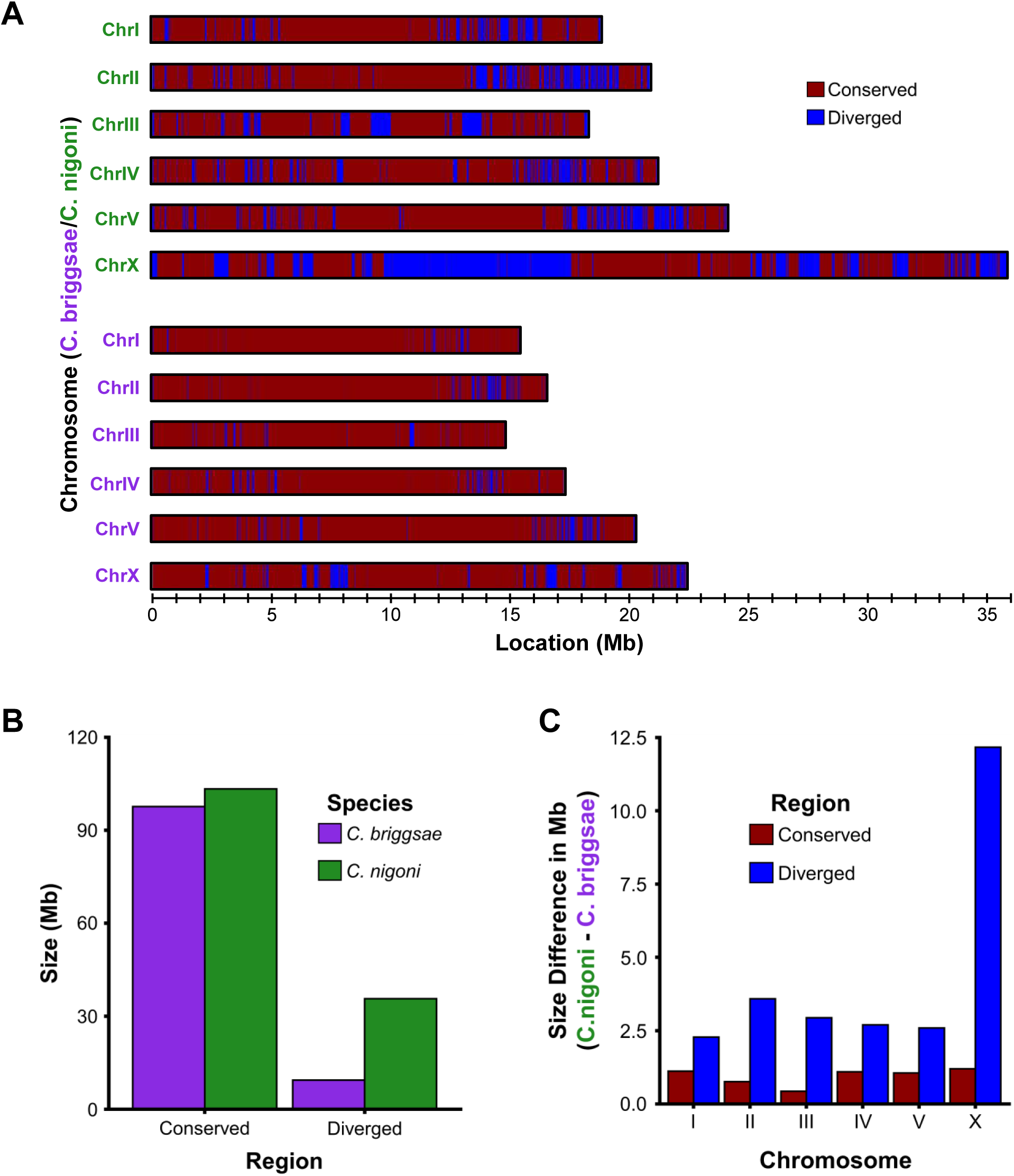
Synteny analysis between the *C. briggsae* and *C. nigoni* genomes. (**A**) Chromatogram demarcating the *C. briggsae* and *C. nigoni* into conserved and diverged regions using Winnowmap2. Red represents conserved regions between the *C. briggsae* and *C. nigoni* genomes, whereas blue represents diverged regions (minimum size of both regions is 10kb). The color of the labels (purple for *C. briggsae* and green *C. nigoni*) denotes the species. Note the preferential location of diverged regions on the distal thirds (“arms”) of the autosomes, as well as the present of large blocks of diverged sequences, especially on the X chromosome. (**B**) Bar graph comparing the cumulative sizes of the conserved and diverged regions between *C. briggsae* and *C. nigoni*. While the majority of both genomes is conserved, the diverged regions are substantially larger in *C. nigoni*. (**C**) Bar graph comparing the size differences between the conserved (red) and diverged (blue) regions across chromosomes. The net change in size is similar for conserved regions across all chromosomes and smaller than the net change for diverged regions. The net change in the size of the diverged regions on the X chromosome is markedly larger than the autosomes.

Next, we analyzed the frequency of chromosomal rearrangements between *C. briggsae* and *C. nigoni*. Comparisons between *C. briggsae* and *C. elegans* have highlighted a high rate of inversions and a scarcity of translocations (Hillier et al. 2007), a pattern that is also observed across more diverged nematode lineages (Carlton et al. 2022). Comparisons between the more closely related *C. briggsae* and *C. nigoni* using the new genomes backed this trend. Using the globally derived synteny alignment and only focusing on rearrangements greater than 10kb, we identified 122 inversions and 2 translocations. This translates to ∼17.4 inversions per million years, a rate that is over 2.5 times higher than the rate derived by comparing *C. briggsae* and *C. elegans* (∼6.9 inversions per million years) (Hillier et al. 2007). Of the two translocations, one was the 45S rDNA array (see below), while the other is a 15kb intra-chromosomal translocation on the X chromosome. Analysis of the inversions reveals that the autosomes contain between 21 and 25 inversions, while the X chromosome only has five, which is significantly less than expected if inversions occur randomly (*p*-value = 1.71 x 10^-8^) (**Figure 4A**). Additionally, the inversions preferentially occur at the arms of chromosome relative to the center (*p*-value = 2.63 x 10^-11^) (**Figure 4B**).

**Figure 4:**
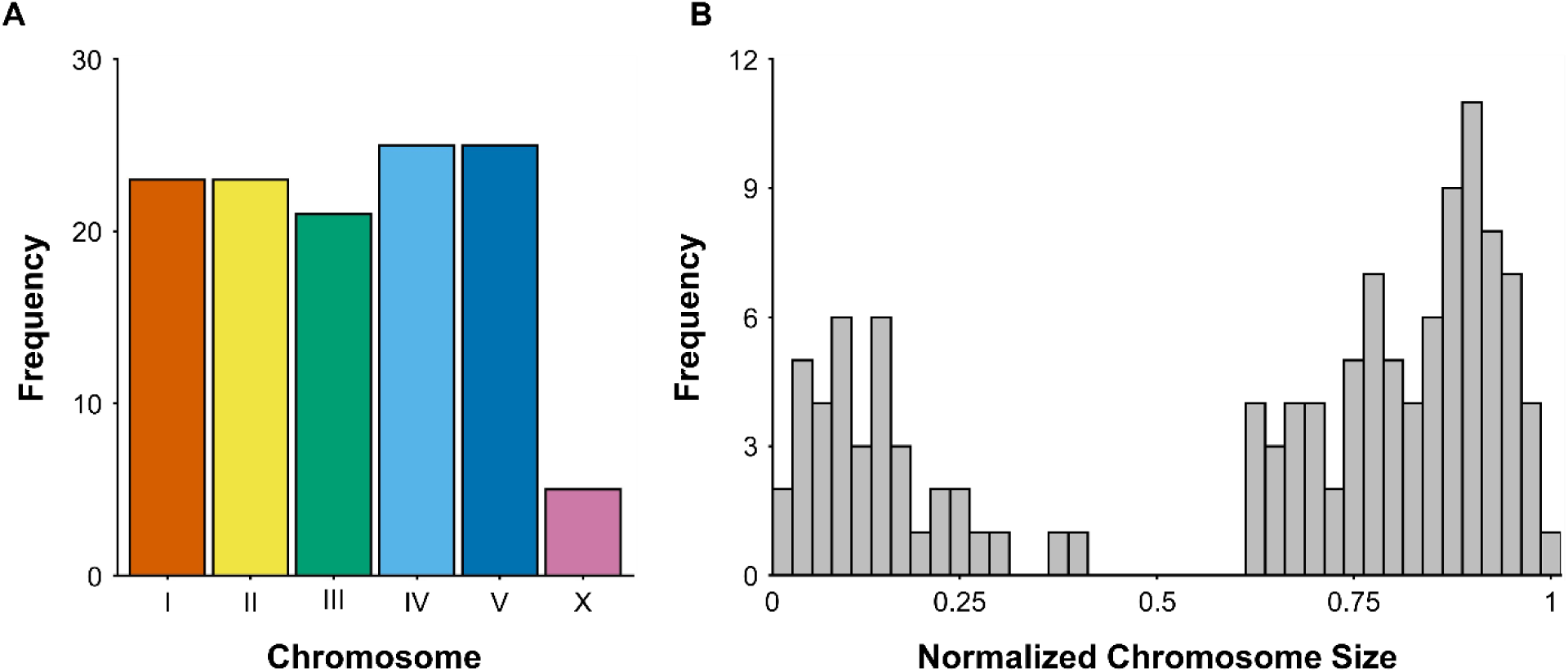
Inversion frequency and distribution between *C. briggsae* and *C. nigoni*. (**A**) Bar graph showing the number of inversions >10kb on each chromosome. The X chromosome has significantly less inversions than expected if inversions occur randomly (*p*-value = 1.71 x 10^-8^, bootstrapping). (**B**) Histogram displays the distribution of all inversions >10kb along a normalized chromosome. Inversions preferentially occur at the chromosome arms relative to the center (*p*-value = 2.63 x 10^-11^, bootstrapping).

Finally, we address the specific genetic features that account for the 32.0Mb difference in genome size between *C. briggsae* and *C. nigoni*. Previous work showed *C. briggsae* has lost many small and male-biased genes that are present in *C. nigoni* (Yin et al. 2018). However, these losses contribute very little to the overall genome size differences (Adams et al. 2023). Differences in introns and exons length similarly have only a small effect on overall genome size (Fierst et al. 2015; Yin et al. 2018). Unlike comparisons of the *C. elegans* and *C. inopinata* genomes (more distantly related androdiecious/gonochoristic sister species), transposable element content does not contribute to the genome size differences observed between *C. briggsae* and *C. nigoni* (Fierst et al. 2015; Kanzaki et al. 2018; Stevens et al. 2019; Adams et al. 2023). We confirmed this observation using our new T2T reference genomes by applying RepeatModeler2 (Flynn et al. 2020) to identify TEs (**Figure 5A and 5B**) (**Supplemental Table 5**). In alignment with the previous studies, we find that *C. briggsae* and *C. nigoni* have similar TE abundance, with TEs accounting for a slightly higher percentage of the genome in *C. briggsae* (8.2% and 7.3% of the respective).

**Figure 5:**
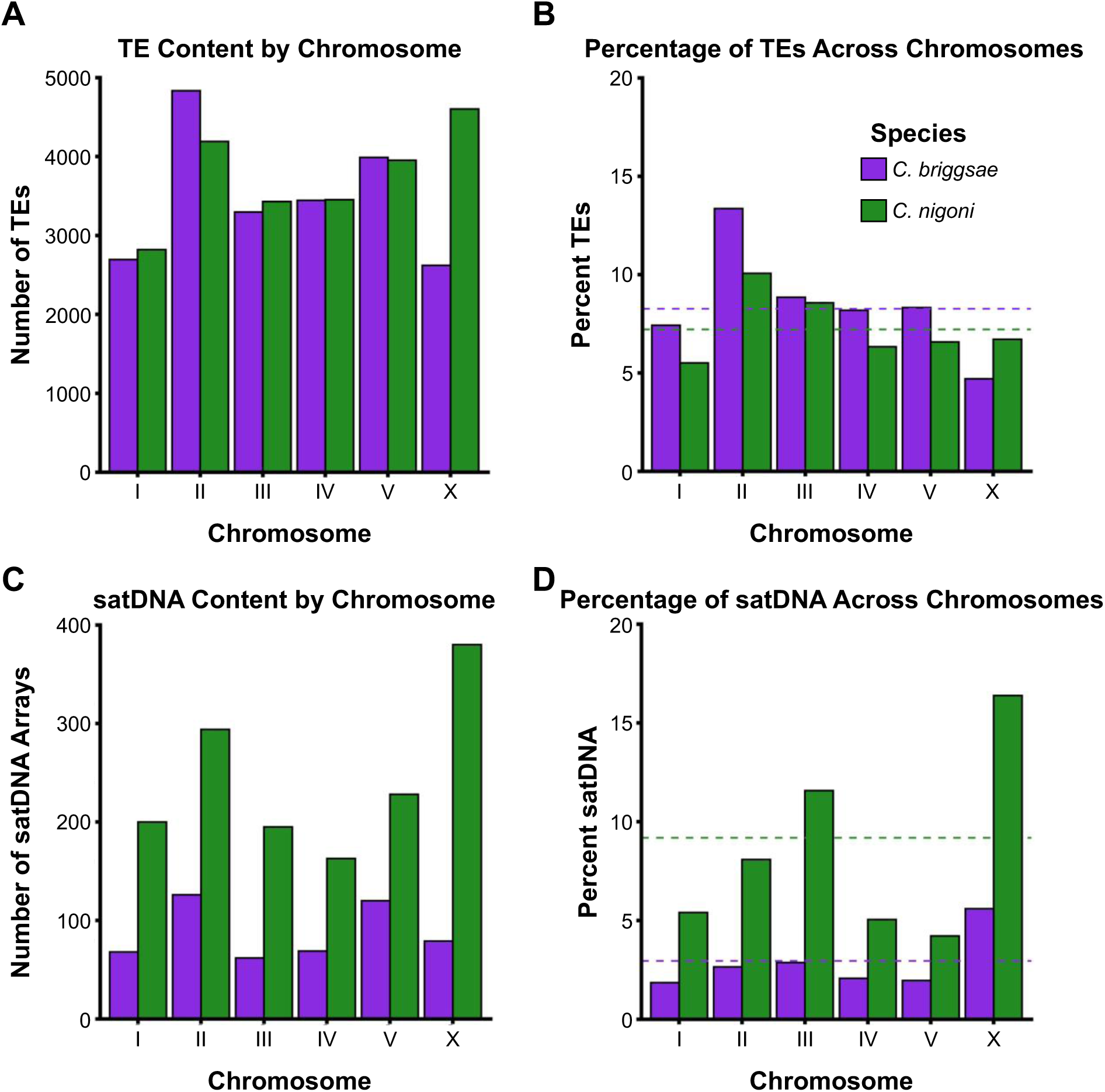
*C. briggsae* and *C. nigoni* TE and satDNA content. (**A**,**C**) Bar graphs showing the number of TEs (**A**) and satDNA arrays (**C**) on each chromosome. (**B**,**D**) Bar graphs showing the percentage of each chromosome occupied by TEs (**B**) or satDNA (**D**). Dashed lines represent the genome-wide fraction. For all graphs, purple denotes *C. briggsae* while green denotes *C. nigoni*. Note the similar per-chromosome TE fraction and the large disparity in per-chromosome satDNA content.

We annotated satDNA using Tandem Repeats Finder (Benson 1999) to determine whether satDNA content explains the differences in genome size. Here, we define satDNA as arrays >1kb with a repeated unit <2kb; this analysis includes most interstitial satDNA and telomeres, but excludes the 45S rDNA and some smaller satDNA arrays, especially microsatellites; note that nematode chromosomes are holocentric and lack centromeric satDNA. Strikingly, we find that *C. briggsae* has roughly a third of the number of satDNA arrays present in *C. nigoni* genome and less than a fourth of the total base pairs in satDNA arrays, reflecting a loss of 9.6Mb (**Figure 5C and 5D**) (**Supplemental Table 6**). satDNA accounts for over 30.0% of the sequences lost in *C. briggsae* despite representing only 9.2% of the *C. nigoni* genome. As expected from the disproportionate loss of sequence on the X chromosome, we find that 4.6Mb of the lost sequences (48.0%) are on the X chromosome.

In summary, we find a much larger degree of genome shrinkage in *C. briggsae* than previously appreciated, with most of the shrinkage occurring on the X chromosome. The vast majority of these lost sequences originate in highly diverged regions where satDNA was disproportionately lost.

### The *C. nigoni* genome contains a second 45S rDNA array

The 45S rDNA arrays persist as one of the greatest challenges preventing the assembly of T2T genomes due to their sheer size, head-to-tail orientation, and sequence homogeneity. In the T2T mouse and human genomes, the 45S rDNA arrays were not assembled via sequencing, but rather artificially arranged based on estimates of sequence abundance and cytology (Nurk et al. 2022; Liu et al. 2024; Potapova et al. 2025). In the recently published *C. elegans* T2T genome, the presence and distribution of single-nucleotide variants (SNVs) allowed for the direct assembly of its lone, 107 copy, 45S rDNA array (Ichikawa et al. 2025).

We similarly assembled or reconstructed the 45S rDNA arrays of *C. briggsae* and *C. nigoni*. Previous work has estimated the *C. briggsae* 45S rDNA array is composed of 85 full copies (Ding et al. 2022). In our assembly, we were able to leverage five SNVs to assemble 68 copies (**Figure 6A**). Based on the previous copy number estimate and absence of any additional SNVs, we reconstructed the remaining 17 copies using the consensus sequence. The lower SNV frequency in the *C. briggsae* 45S rDNA relative to the *C. elegans* N2 array (average 1 SNV every 128kb and 23kb, respectively) accounts for the need to reconstruct this last section. In total, this places the *C. briggsae* 45S rDNA array size at ∼640kb on the left end of chromosome V.

**Figure 6:**
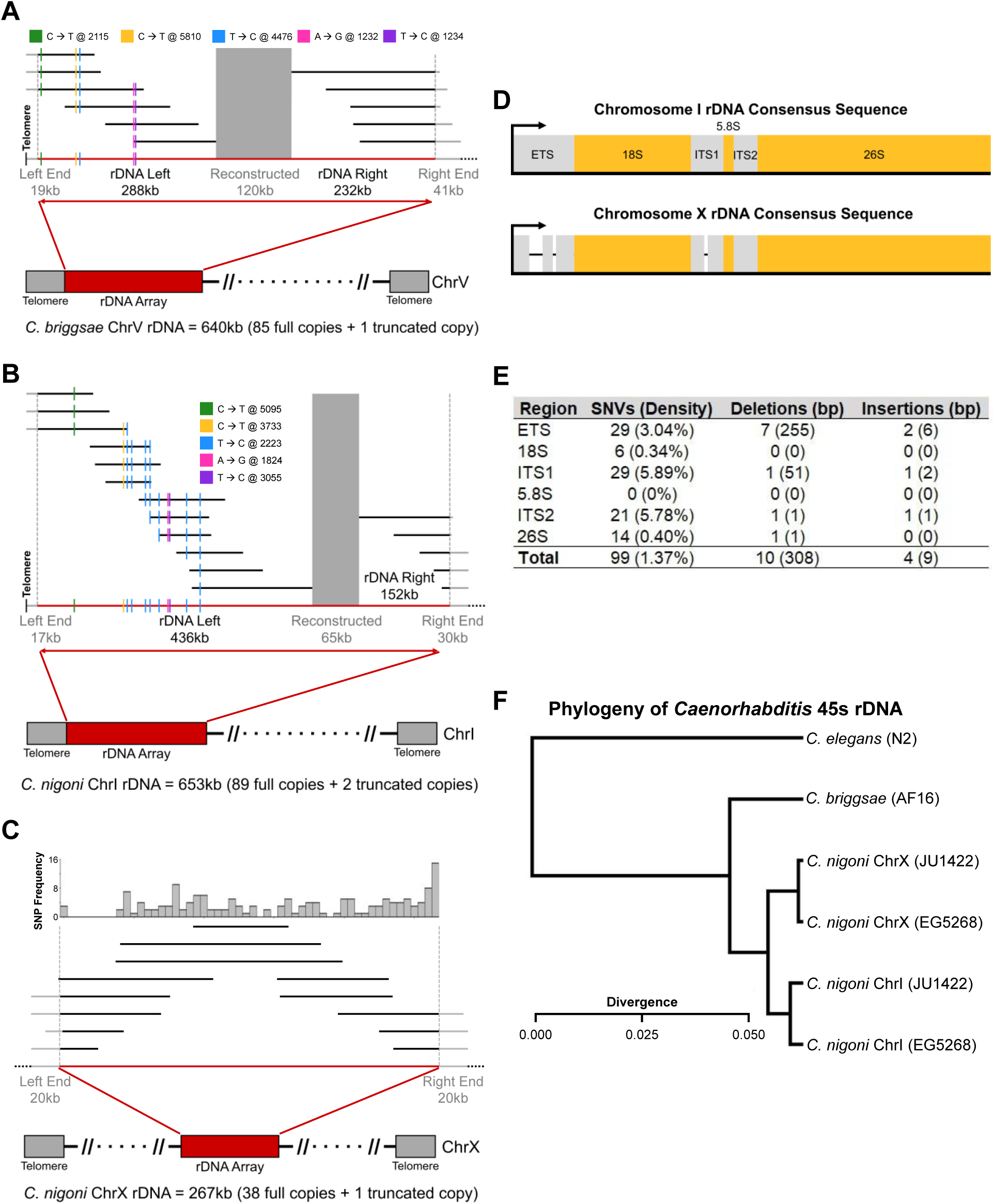
45S rDNA arrays of *C. briggsae* and *C. nigoni*. (**A-C**) Assembly and reconstruction of the 45S rDNA arrays for *C. briggsae* chromosome V (**A**), *C. nigoni* chromosome I (**B**), and *C. nigoni* chromosome X (**C**). Black horizonal lines highlight a subset of Nanopore reads (to scale), aligned to the 45S rDNA arrays, that were used to assemble the array. Grey portions on the arms on some of the black horizontal lines represent anchor regions that align to the unique sequences that flank the 45S rDNA arrays. In **A** and **B**, vertical-colored lines represent single-nucleotide variants (SNVs) within the consensus 45S rDNA sequence for the given array, while the large grey boxes highlight the reconstructed region. Schematics below the Nanopore reads depict the genetic structure of the arrays. In **C**, due to the high density of SNVs within the array, the distribution of SNVs is represented by a histogram located above the Nanopore reads. (**D**) Schematic diagrams comparing the genetic structures of the *C. nigoni* chromosome I and chromosome X 45S rDNA consensus sequences. The 5.8S, 18S, and 26S coding regions are yellow, while the external transcribed spacer (ETS) and the internal transcribed spacers (ITS1 and ITS2) are grey. Deletions over 50bp that are present in the chromosome X 45S rDNA consensus sequence are denoted by black dashes. (**E**) Table showing the frequency, by region, of SNVs, deletions, and insertions in the *C. nigoni* chromosome X 45S rDNA consensus sequence relative to the *C. nigoni* chromosome I 45S rDNA consensus sequence. Note the very low frequency of variants in the 18S, 5.8S and 26S regions. (**F**) Phylogenetic tree of the 45S rDNA consensus sequences for *C. elegans* (from (Ellis et al. 1986)), *C. briggsae* and *C. nigoni* (strains JU1422 and EG5268), showing that the *C. nigoni* chromosome I and X chromosome 45S rDNA arrays from the two diverged isolates cluster together.

During our assembly of the *C. nigoni* genome we identified two 45S rDNA arrays, one on chromosome I and another on the X chromosome. Of the two arrays, the X chromosome array was easier to assemble because of a high density of SNVs (average of 1 every 1.6kb) and maximum distance between SNVs of only 40kb. (**Figure 6B**). Thus, we were able to assemble the 39 copies that make up the 267kb array from sequencing alone. We then used the coverage of this array to estimate the size of the chromosome I array (similar to (Ding et al. 2022), see **Methods**) to be 89 copies. Of the 89 copies, we were able to use 11 SNVs (average 1 SNV every 59kb) to assemble 80 copies, with the remaining nine copies reconstructed using the consensus sequence, making the total size of the array ∼653kb (**Figure 6C**).

Although the presence of 45S rDNA arrays on multiple chromosomes is common across the animal kingdom, to date, no nematode has been shown to possess multiple arrays (Sochorova et al. 2018; Hall et al. 2022). Comparing the consensus 45S rDNA unit from both arrays revealed that each array is composed of different variants. Specifically, the variant present on left arm of chromosome I corresponds to the 45S rDNA units present on the left arm of chromosome V in *C. briggsae* and the right arm of chromosome I in *C. elegans*, suggesting it reflects the ancestral 45S rDNA array that translocated between the chromosomes. The variant on the X chromosome contains three large deletions (>50bp) within the external transcribed spacer (ETS) and the internal transcribed spacer 1 (ITS1) (**Figure 6D**). The distribution of these large deletions as well as of smaller indels and SNVs, which are rare within 5.8S, 18S, and 26S structural ribosome subunits (**Figure 6E**), suggest that the X chromosome array is functional.

Finally, we analyzed rDNA sequences in the highly diverged *C. nigoni* EG5268 strain (Kiontke et al. 2011). The phylogeny of the consensus *C. nigoni* 45S rDNA unit revealed that they cluster by chromosome and that the arrays are more similar to each other than to the *C. briggsae* 45S rDNA cluster (**Figure 6F**). This finding supports the idea that this second 45S rDNA array is functional and suggests that its existence is fixed in *C. nigoni*.

## Discussion

Here, we assembled complete, gapless T2T genomes of *C. nigoni* and *C. briggsae* and used them to deduce the evolutionary dynamics between these closely related species. Our results provide insight into the genome contraction events that coincided with the transition of *C. briggsae* to a self-fertilizing reproductive mode. We identify satDNA as a genomic feature that was disproportionally lost during the rapid contraction of the genome. We also found many inversions between *C. nigoni* and *C. briggsae*, as well as very few interchromosomal rearrangements, consistent with the high degree of karyotype conservation across the *Caenorhabditis* genus. Using our substantially improved *C. nigoni* reference genome, we identified a second, likely functional 45S rDNA array on the X chromosome. The resources generated in this study will undoubtedly improve the scope of future genomic studies involving the *Caenorhabditis* genus, an established collection of species that will continue to provide wide-ranging insights across the many fields of biology.

While previous work documented genome shrinkage associated with the transition to self-fertilization, our work contributes to our understanding of both the magnitude of shrinkage and the mechanisms underlying it. By comparing gapless T2T genomes, we deduced that the *C. briggsae* genome is 32Mb smaller than *C. nigoni*, compared with a 21Mb difference calculated based on previous genome assemblies (**Table 2**). Many evolutionary factors including effective population size, effective recombination, mode of reproduction, mutation rates, natural selection, transmission distortion, and TE bursts contribute to the evolution of genome size. As a result, two main models, the mutational hazard (Lynch and Conery 2003) and the mutation equilibrium (Petrov 2002), are primarily used to describe trends in genome size variation. The mutational hazard model is based on the premise that a large effective population size entails high selection efficacy, which results in a small and streamlined genome. The reciprocal is that a small effective population size prevents efficient removal of weakly deleterious mutations, resulting in a larger genome. *Caenorhabditis* species exhibit the opposite trend. The self-fertilizing *Caenorhabditis* species *C. briggsae* (Sivasundar and Hey 2003), *C. elegans* (Dolgin et al. 2008), and *C. tropicalis* (Noble et al. 2021) have an effective population size of around 10^4^, while the outcrossing species *C. brenneri* (Dey et al. 2013; Teterina et al. 2025), *C. remanei* (Graustein et al. 2002; Cutter et al. 2006; Teterina et al. 2023), and presumably *C. nigoni* (Teterina et al. 2025) have effective population sizes that are orders of magnitude greater (10^5^ - 10^6^). However, self-fertilizing species have smaller genomes than their outcrossing species, with the *C. briggsae* genome 29.9% smaller than *C. nigoni*.

Alternatively, the mutational equilibrium model conceptualizes genome size as the interplay between insertions and deletions. Within this, the accordion model (Kapusta et al. 2017) postulates that genome size expansions and contractions can occur rapidly due to the interplay between large duplications or TE bursts on one hand, and large deletions on the other (Kapusta et al. 2017). As with previous studies, we identified no changes in abundance of TEs between *C. briggsae* and *C. nigoni* (**Figure 5**) (Fierst et al. 2015; Adams et al. 2023). This is consistent with simulations that demonstrate that the chances of a TE invading a genome is drastically reduced when self-fertilization reduces the effective population size (Hua-Van et al. 2011). Additionally, the rare occurrence of males is likely sufficient to prevent TE accumulation via Muller’s ratchet, despite largely reduced effective recombination in self-fertilizing species (Muller 1964; Dolgin and Charlesworth 2008). While we find no evidence for large duplications in the lineage leading to *C. nigoni*, our data supports an important role for large deletions in *C. briggsae*. Our synteny analysis revealed that the genome shrinkage of *C. briggsae* was driven by large deletions that were disproportionately satDNA (**Figure 3 and 5**). Dynamic in nature, satDNA mutates at a rate 10-100,000 times greater than the rest of the genome (Gymrek et al. 2017). At least in some contexts, there is also experimental evidence that suggest that, in *C. briggsae* specifically, deletions of satDNA are more prevalent than insertions (Phillips et al. 2009b). Overall, our data is well aligned with the accordion model. We propose that genome shrinkage in *C. briggsae* is likely due to a preponderance of large deletions. The transition to self-fertilization drastically lowered the rate of effective recombination, leading to a sharp decrease in effective population size, which, in turn, allowed for genetic drift to quickly fix such deletions.

Our revised comparative analysis also revealed a previously unappreciated disparity in sequence loss between the X chromosome and the autosomes. Specifically, we found that the X chromosome is responsible for 41.8% of the genome shrinkage (**Table 2 and Figure 2**). While a wider survey of *Caenorhabditis* could reveal whether this is a recurring feature, the X chromosome is distinct in many aspects (Cutter 2018). For example, the X chromosome replicates later than the autosomes during meiotic S phase (likely reflecting it unique chromatin state) (Jaramillo-Lambert et al. 2007) and it possesses a relatively smaller chromosome center (based on its recombination landscape) (Rockman and Kruglyak 2009; Ross et al. 2011). These and other X-specific features may contribute to the differences we observed here. For instance, shorter autosomes in *Caenorhabditis* preferentially segregate with the lone X chromosome in male meiosis. Given the large contribution of rare males to gene flow in androdiecious species, such transmission distortion bias could rapidly shrink the autosomes following the transition to self-fertilization (Wang et al. 2010; Fierst et al. 2015; Le et al. 2017). However, this bias could not explain the even faster shrinkage of the X chromosome we observed in *C. briggsae*. One mechanism that could underlie the disproportionate loss of chromosome X sequence is its smaller effective population size relative to the autosomes - due to the XO/XX sex determination system - at the time of the transition. Nevertheless, it is likely that additional mechanisms contribute to preferential sequence loss on the X chromosome.

Through our analysis of genome synteny between *C. briggsae* and *C. nigoni*, we identified that the X chromosome possessed significantly less inversions than the autosomes (**Figure 4**). This is expected because *C. nigoni* is an outcrossing species so the hemizygosity of the X chromosome in males exposes deleterious alleles, which consequently reduces the likelihood of deleterious, polymorphic inversions from persisting (Connallon et al. 2018). Additionally, this hemizygosity translates to the effective population size the X chromosome being smaller, which means that genetic drift is stronger. Here, we estimated that the rate of inversions to be roughly two and a half times higher than the rate derived by comparing *C. briggsae* to *C. elegans* (Hillier et al. 2007). This increased measured rate of inversions could be due to the relatively short evolutionary time since the species split, which allows us to precisely observe all inversions. However, we cannot rule out idiosyncratic effects on inversions in different lineages.

We documented a surprising degree of plasticity in the genomic location of the 45S rDNA arrays, with the location of the ’main’ array changing between *C. elegans*, *C. briggsae* and *C. nigoni*, and the *C. nigoni* genome containing a second array on the X chromosome (**Figure 6**). While it is common for animal genomes to have multiple 45S rDNA arrays across multiple chromosomes with each chromosome sometimes having a specific variant (Hall et al. 2022), it has not been previously observed in *Caenorhabditis*. Here, we annotated its presence and compared its sequence homology to the 45S rDNA present in *C. briggsae*, *C. elegans*, and *C. nigoni*. Our analysis suggests that the second, X chromosome array is involved in ribosome biogenesis based on it mutation patterns and its conservation. In addition, rDNA array can also play functional roles in genome organization, genome stability, and the regulation of replication and transcription (Kobayashi 2008; Yu and Lemos 2016; Kwan et al. 2023). Humans, mice, and plants have different 45S rDNA variants with tissue-specific expression patterns. The accessibility of the rDNA arrays of *Caenorhabditis* to long-read sequencing, specifically this newly discovered array in *C. nigoni* which was completely assembled (rather than reconstructed), offers an exciting system to probe its biological functions and investigate the consequences of its perturbation.

Advances in third-generation sequencing enable the construction of T2T reference genomes from sequencing alone. The recent, gapless T2T assembly of the *C. elegans* reference genome provided key proof of concept and became the first animal model organism to have a T2T genome assembled from a wild isolate (Ichikawa et al. 2025). This is distinct from the recent mouse (Liu et al. 2024) and human (Nurk et al. 2022) T2T genome assemblies, which contain regions that could only be reconstructed using imaging and computational approaches, and that are based on a haploid cell line (mhaESC) and a homozygous complete hydatidiform mole (CHM13), respectively (Potapova et al. 2025). Notably, the *C. elegans* genome was assembled via a combination of high-depth sequencing from Nanopore and PacBio platforms (287x and 303x coverage, respectively). Here, we used ∼40x sequencing coverage (∼5Gb) of Nanopore reads to generate two T2T reference genomes. For perspective, typical yields for Nanopore MinION and PromethION flow cells are ∼15Gb and ∼100Gb, respectively, meaning that roughly three and ∼20 T2T genomes can be generated from a single run of each. These advances render *Caenorhabditis* species a promising model system for comparative- and population-genomic studies that consider the complete repertoire of genomic elements.

## Materials and Methods

### Worm maintenance and DNA extraction

*C. briggsae* and *C. nigoni* strains were cultured as previously described (Sulston and Brenner 1974). In brief, worms were grown on 15cm 2XYT media plates seeded with NA22 *E. coli* until they starved the plate (Sambrook 1989). Starved worms were rinsed with wash buffer (100mM Tris-HCl pH 8.0, 100mM NaCl, 20mM EDTA) three times. The second wash included a 30-minute incubation at room temperature to help clear remaining bacteria. Worms were then pelleted and 50µL of worm slurry was resuspended in 485µL of wash buffer and frozen at −80°C for a minimum of 10 minutes (Tyson et al. 2018). The worms were then thawed and digested using Proteinase K (0.4mg/mL final concentration) and SDS (0.2% final concentration) for 2 hours at 60°C. Following this initial digestion, 500µL of lysis buffer (100mM Tris-HCl pH 8.0, 100mM NaCl, 20mM EDTA, 4mg/mL Proteinase K and 0.2% SDS) was added and incubated overnight. RNAse (2µg/mL final concentration) was added and incubated at 37°C for 30 minutes. DNA was extracted by performing a phenol-chloroform extraction and ethanol precipitation (Ashburner 1989; Tyson 2020; Kim et al. 2021). DNA quality and quantity was checked using a NanoDrop spectrophotometer and a Thermo Fisher Scientific Qubit 4 Fluorometer.

### Nanopore sequencing

Oxford Nanopore Technologies (ONT) libraries were prepared using ONT SQK-LSK114 according to (Tyson 2020; Kim et al. 2021). Samples were sequenced to at least 40x coverage (>5Gb) using an ONT MinION sequencing device (Tyson 2020; Kim et al. 2021). All basecalling was performed using the ONT MinKNOW software at the time of sequencing and the super-accurate basecalling model (for versions, see header of fastq files), with only passed reads and reads over 1kb being kept for downstream analysis. Sequencing quality was analyzed using NanoPlot (v.1.41.3) (De Coster and Rademakers 2023).

### Genome assembly and annotation

Initial draft assemblies were done by performing *de novo* genome assembly using Flye (v.2.9.1-b1780) (Kolmogorov et al. 2019) with the following parameters; --genome-size 150m --threads 56 --iterations 5 --min-overlap 10000 --asm-coverage 40. Initial chromosome assignment was determined by mapping the contigs to previous *C. briggsae* and *C. nigoni* reference genomes (Yin et al. 2018; Stevens et al. 2022). To ensure quality and completeness, each assembly was manually inspected using IGV (v.2.16.1) (Robinson et al. 2023) (**Supplemental Figure 1**). If a gap persisted between two contigs, reads greater than 50kb in length were mapped to the contigs using Winnowmap2 (v.2.03) (Jain et al. 2022) with the following additional parameter; -k 28. The soft clippings of these reads were used to extend the contigs until the two contigs were bridged. After the contigs were bridged, the subsequent sequence was polished with Flye (-- iterations 5) and manually inspected. This method for elongating contigs was similarly employed by (Ichikawa et al. 2025). In other instances, Flye would misassemble a region (**Supplemental Figure 2**). In these cases, the region would be manually corrected and polished using Flye. In total, the *C. briggsae* assembly had 21 instances where contigs need to be assembled and 17 issues where Fyle misassembled a region, while the *C. nigoni* assembly had 32 instances where contigs need to be assembled and 20 issues where Fyle misassembled a region. Another issue stemmed from strand-specific basecalling errors by the Nanopore basecaller (**Supplemental Figure 3**). These errors occurred 44 and 14 times in *C. briggsae* and *C. nigoni*, respectively, and were resolved by polishing with either PacBio (PRJNA384657) or Illumina reads (PRJNA232381) (Albritton et al. 2014; Yin et al. 2018). Two regions that required additional assistance were the telomeres and the 45S rDNA arrays (see below). While the *C. nigoni* JU1422 was almost completely homozygous, we identified a 10.6 Mb region of residual heterozygosity on chromosome III which was manually phased using Flye, IGV, and Winnowmap2. This residual heterozygosity was also present in the PacBio reads used in the prior assembly (Yin et al. 2018). We included only one of the two haplotypes in the assembly, while the second was separately added to GenBank (PRJNA1418286).

Telomere length is highly variable depending on many factors including cell type, cell division history, telomerase activity and environmental factors (Greider 1996). The consensus across *Caenorhabditis* is that telomere length is typically 2-9kb (Raices et al. 2005; Frenk et al. 2019), making their assembly accessible using our Nanopore reads. Here, telomere length was determined by the read with the longest telomere length that still anchored to the subtelomere region (**Supplemental Figure 4**). In all cases, additional reads terminated within ∼1kb of the longest read. This resulted in telomere lengths ranging from 3.5kb to 7kb in *C. brigssae* (median = 5,703bp) and 800bp to 3kb in *C. nigoni* (median = 1,722bp).

The *C. nigoni* chromosome X 45S rDNA array had a high enough degree of structural variations that it could be assembled by elongation method mentioned above using Flye, Winnowmap2, and IGV. The *C. briggsae* chromosome V 45S rDNA array and the *C. nigoni* chromosome I 45S rDNA array were assembled by identifying SNVs within the rDNA units (**Figure 6A-C**). First, we identified all reads greater than 50kb and containing a complete rDNA unit and classified them based on whether they contained an anchor sequence. An anchor sequence was defined as the unique non-rDNA sequences that flank the rDNA arrays. SNVs were identified by elongating from the anchor regions and also by identifying SNVs that were at a greater frequency than the expected noise. More specifically for the latter, SNVs were identified by extracting all complete rDNA units from Nanopore reads and aligning them to a previously curated *C. briggsae* 45S rDNA unit (Ding et al. 2022) and polished with Flye to generate consensus sequences for both the new *C. briggsae* genome and *C. nigoni* genome.

The 45S rDNA units were then aligned to the consensus sequence and SNVs were identified using IGV tools. SNVs were filtered based on frequency and whether they occurred on both strands (i.e., strand-specific SNVs were filtered out). Candidate SNVs had to have a minimum coverage of 10 for *C. nigoni* and *C. briggsae*, respectively. For context, the *C. briggsae* AF16 45S rDNA array was estimated to contain 85 full copies (Ding et al. 2022), while the filtered set Nanopore reads producing 1,661 complete rDNA units (∼19.5x coverage). Based on alignment to the reference, the alignment error rate was 0.5% (65,991 errors / 12.4Gb), meaning that each nucleotide that did not match the consensus nucleotide had an average frequency of 2.6 errors (0.5% alignment error rate x 1,661 coverage / 3 possible incorrect nucleotides). Note that a true SNV would be around the coverage and substantially greater than this error. By combining the two approaches, we were able to mostly assemble the 45S rDNA arrays, by identifying reads that contained multiple SNVs. The 45S rDNA copy number for *C. nigoni* chromosome I was estimated by taking the number of 45S rDNA units aligning to chromosome I (1,272), dividing it by the number of 45S rDNA units aligning to chromosome X (417), multiplying it by ratio of chromosome X to chromosome I (males are XO and females XX, so the ratio is 0.75) and multiplying it by the number of 45S rDNA copy number for the directly-assembled *C. nigoni* chromosome X (39), which produces the estimated copy number of 89. The phylogenetic tree of the consensus 45S rDNA unit was generated using MEGA12 (v.12.0.9) (Kumar et al. 2024) (parameters: statistical method = UPGMA, Test of Phylogeny = Bootstrap Method, No. of Bootstrap Replications = 1000, Model/Method = Kimura 2-para, and Gaps/Missing Data Treatment = Complete Deletion) and plotted using phytools (v2.4-4) (Revell 2012).

With the rDNA arrays complete, genome quality was assessed by calculating BUSCO (Simao et al. 2015) completeness scores using compleasm (Huang and Li 2023), which checks for expected single-copy genes. Protein-coding genes were predicted using Augustus (Stanke et al. 2006). Chromatograms were generated by using Winnowmap2 to align the old reference genomes to these new reference genomes and plotted using karyoploteR (v.1.32.0) (Gel and Serra 2017). Transposons were annotated using RepeatModeler2 (Flynn et al. 2020) as implemented by reasonaTE (Riehl et al. 2022). satDNA was defined as any tandem repeat where the motif repeated at least five times and the length of the array was greater than 1kb. satDNA were annotated using Tandem Repeats Finder (Benson 1999) (parameters = matching weight = 2, mismatching penalty = 7, indel penalty = 7, match probability = 80, indel probability = 10, minimum alignment score to report = 50, maximum period size to report = 2000). Based on limitations in the Tandem Repeats Finder algorithm, only satDNA with period sizes of 2000bp or less were considered. For example, satDNA arrays like the 45S rDNA arrays were not called by Tandem Repeats Finder and thus were not considered satDNA for analysis purposes.

### Immunofluorescence

Immunofluorescence was performed as in (Phillips et al. 2009a). In brief, worms were grown on NGM plates seeded with OP50 (Stiernagle 2006). L4 worms (hermaphrodites of *C. briggsae* and a combination of males and females for *C. nigoni*) were picked and transferred to fresh plates and grown for 48 hours. *C. briggsae* hermaphrodites and *C. nigoni* females were placed onto a coverslip with 1x EBT (1.1x Egg Buffer, 0.1% Tween-20, 0.0065% tetramisole). Gonads were then dissected by cutting off the heads and tails with a scalpel before adding 1x Fix (1x Egg Buffer, 2% formaldehyde) in 1:1 ratio with the 1x EBT. Samples were mixed by pipetting. Excess liquid was removed, and the samples and transferred to Histobond slides where they were then frozen on dry ice. Coverslips were then freeze-cracked off of the slide and samples were fixed in methanol. Gonads were then rinsed three times in 1x PBST, incubated for 30 minutes in Roche Block, and incubated overnight in rabbit anti-H3K36me3 (1:1,000; Active Motif #61102) primary antibody at 4°C in a humid chamber. Samples were then washed with 1x PBST three times before incubating in donkey anti-rabbit (1:500; Jackson ImmunoResearch #711-605-152) secondary antibody for two hours at room temperature in a humid chamber. Finally, samples were washed in 1x PBST, stained with DAPI (1:10,000) in 1x PBST, washed with 1x PBST and mounted in mounting medium (70mM TRIS, 45% NPG-Glycerol).

### Image acquisition and chromosome tracing

Gonads were imaged using the 63x 1.4 NA oil immersion objective on a Zeiss LSM880 confocal microscope in AiryScan mode. Images were processed using Zen Blue 3.6 (Zeiss) software by performing Airyscan processing, followed by Channel Alignment (parameters: Preprocessing = No Preprocessing, Quality = Highest, Registration Method = Translation, Interpolation = Cubic, Third Dimension = Z). Chromosome tracings were performed using Imaris (version 10.2.0, Oxford Instruments). Specifically, the Surfaces function was applied individual nuclei using Absolute Intensity Thresholding. In some cases, Surfaces clustered the X chromosome with the autosomes. If this occurred, the parameter “split touching objects” was applied and the autosome and X chromosome clusters were manually assigned. Finally, area, volume, and intensity sum statistics were calculated for each cluster and reported as the statistic of the X chromosome divided by the sum of the X chromosome and the autosomes (**Supplemental Table 3**). This percentage of total was used because area and volume vary by pachytene stage and intensity sum varies by sample.

### Genome synteny analysis

Genome homology and synteny between the T2T *C. briggsae* and *C. nigoni* genomes were calculated using BLAST (Camacho et al. 2009) and WinnowMap2 (Jain et al. 2022). For BLAST, each *C. nigoni* chromosome was fragmented 10kb non-overlapping regions and blasted against the homologous *C. briggsae* chromosome using blastn with default parameters. Importantly, this method allows for the alignment of every 10kb region independent of the neighboring region, which makes it robust for the identification of large duplications. WinnowMap2 was used to align the complete genomes of *C. briggsae* and *C. nigoni* to each other. To decrease the number of supplemental mappings, the *C. briggsae* genome was fragmented into 500kb non-overlapping regions before being aligned to the *C. nigoni* genome. Conserved regions were defined as regions that were able to align with at least 10kb. Given that WinnowMap2 aligned the complete *C. briggsae* genome to the complete *C. nigoni* genome, we were able to identify inversions and translocations over 10kb. We also conducted the inverse comparisons using both approaches.

To determine if the number of inversions that occur on the autosomes and chromosome arms are more than expected, a set of random inversions (equal in size to the number of observed inversions) was generated. The average number of inversions that occurred on the X chromosome, as well as whether the inversions occur at the chromosome centers (middle 50%) or on the arms (each 25%) was calculated. This simulation was bootstrapped 10,000 times to build a distribution, where the observed value could be compared and a *p*-value could be derived. Pearson Correlation Coefficient was calculated between the two methods of synteny by using the cor.test() function in R by comparing the binary assignments of WinnowMap2 to the percent identity of BLAST.

### Data Availability

All raw sequencing reads and genome assemblies generated in this study are publicly available from the NCBI Sequence Read Archive and GenBank (PRJNA1418286). Reviewer Link: https://dataview.ncbi.nlm.nih.gov/object/PRJNA1418286?reviewer=egpaqgl6ndvt9is85f6hutofdk

## Acknowledgements

We thank the Rog lab, Erik Anderson and Talia Karasov for discussions. Worm strains were provided by the Caenorhabditis Genetics Center, which is funded by NIH Office of Research Infrastructure Programs (P40 OD010440).

